# Crystal structure of the Tof1-Csm3 (Timeless-Tipin) fork protection complex

**DOI:** 10.1101/2020.01.24.918474

**Authors:** Daniel B. Grabarczyk

## Abstract

The Tof1-Csm3 fork protection complex has a central role in the replisome – it promotes the progression of DNA replication forks and protects them when they stall, while also enabling cohesion establishment and checkpoint responses. Here, I present the crystal structure of the Tof1-Csm3 complex from *Chaetomium thermophilum* at 3.1 Å resolution. The structure reveals that Tof1 is an extended alpha-helical repeat protein which is capped at its C-terminal end by Csm3, a small helical bundle protein. I also characterize the DNA binding properties of the complex and a cancer-associated peptide-binding site. This study provides the molecular basis for understanding the functions of the Tof1-Csm3 complex, its human orthologue the Timeless-Tipin complex and additionally the Drosophila circadian rhythm protein Timeless.

## Introduction

As well as error-free DNA synthesis, the eukaryotic replication fork must coordinate processes such as establishing chromosome cohesion, activating the S phase checkpoint and transferring epigenetic material. Furthermore, the polymerases and helicases which form the core of the replication machinery are regulated to couple unwinding and synthesis, protect the fork when it is blocked, and integrate external signals. The coordination of all of these processes is required to maintain genome stability [1]. This becomes especially important in conditions of replication stress, which is a common feature of cancer cells [2, 3]. Indeed many non-essential replisome-associated proteins are upregulated and become essential in cancer [4–6].

The Tof1-Csm3 fork protection complex was identified as a non-essential chromosome cohesion factor [7, 8] that interacts with Topoisomerase 1 [9] to link concatenation and fork regulation [10]. Additionally it has an important role in protecting stalled forks and enabling their restart [11–13], coupling the replicative helicases and polymerases [14], promoting fork progression [15, 16], mediating the S phase checkpoint [17, 18] and maintaining genome stability at CAG repeats [19]. The mammalian orthologue of Tof1-Csm3 is the Timeless-Tipin complex [20], which has similar functions [21–24] and additionally regulates the fork in response to oxidative stress [25]. Timeless also has a PARP1 binding (PAB) domain at its C-terminus important for double-stranded break repair [26, 27], and interacts with RPA through the C-terminus of Tipin [28, 29]. Timeless was first discovered as a circadian rhythm regulator in drosophila [30]. However, drosophila also contain a homologue of mammalian Timeless, known as Tim2, which is likely to be the true orthologue of Tof1 [31]. Nevertheless, it has been suggested that mammalian Timeless links the circadian rhythm with DNA replication [32, 33].

A lack of structural information has precluded progress in the understanding of the specific role that Tof1-Csm3 plays at replication forks, and how this relates to the diverse phenotypes resulting from its mutation. A crystal structure of a the N-terminal domain of Timeless shows that this forms an Armadillo repeat protein [34], while 2D cryo-EM classes of the yeast replisome show that Tof1-Csm3 binds in front of the fork to stabilize incoming DNA [35]. Here, I present the structure of the *Chaetomium thermophilum* Tof1-Csm3 complex. The structure reveals that the protein is folded as a single unit, with Csm3 forming an alpha-helical bundle that caps the Armadillo repeats of Tof1. This suggests a structural role for this complex at the fork. The crystallographic packing in my structure reveals a peptide-binding patch that is affected by a cancer-associated mutation in human Timeless, and I map a double-stranded DNA binding activity to a minimal Tof1-Csm3 complex. The structure also enables sequence alignment of Tof1 with human and drosophila Timeless clarifying the similarity of their structures but differences in function.

## Results

### Structure of the Tof1-Csm3 complex

To determine the structure of the Tof1-Csm3 complex, multiple constructs from the mildly thermophilic fungus *Chaetomium thermophilum* were screened for purification and expression. Crystals were obtained for many of these (Fig. S1A), but only formed in customized crystallization screens designed for challenging complexes. These screens are detailed in Figure S1B, and were also previously used to crystallize another complex [36]. The only construct resulting in diffracting crystals was Tof1(1-728)DL1,2,3-Csm3(48-157). Here, three predicted disordered loops were deleted from Tof1: L1 (256-363), L2 (420-434), L3 (558-585) (Fig. S1A). These crystals diffracted anisotropically to 3.1 Å (Table 1), and the dataset could be solved by molecular replacement using the N-terminal fragment of Timeless (PDB 5MQI) [34] with the remaining half of the protein manually built, exploiting molecular dynamics force-field refinement [37] and contact prediction [38]. A final R_work_/R_free_ of 23.7/26.4% was achieved. The entire structure is well resolved, aside from the N-terminal portion of Csm3, and two small loops in Tof1 (Fig. S2). The main crystal contacts occur at the N-terminus of Tof1, and thus Csm3 and the very C-terminus of Tof1 have relatively high B factors and there is lower map quality in this region (Fig S2C).

**Table 1.**
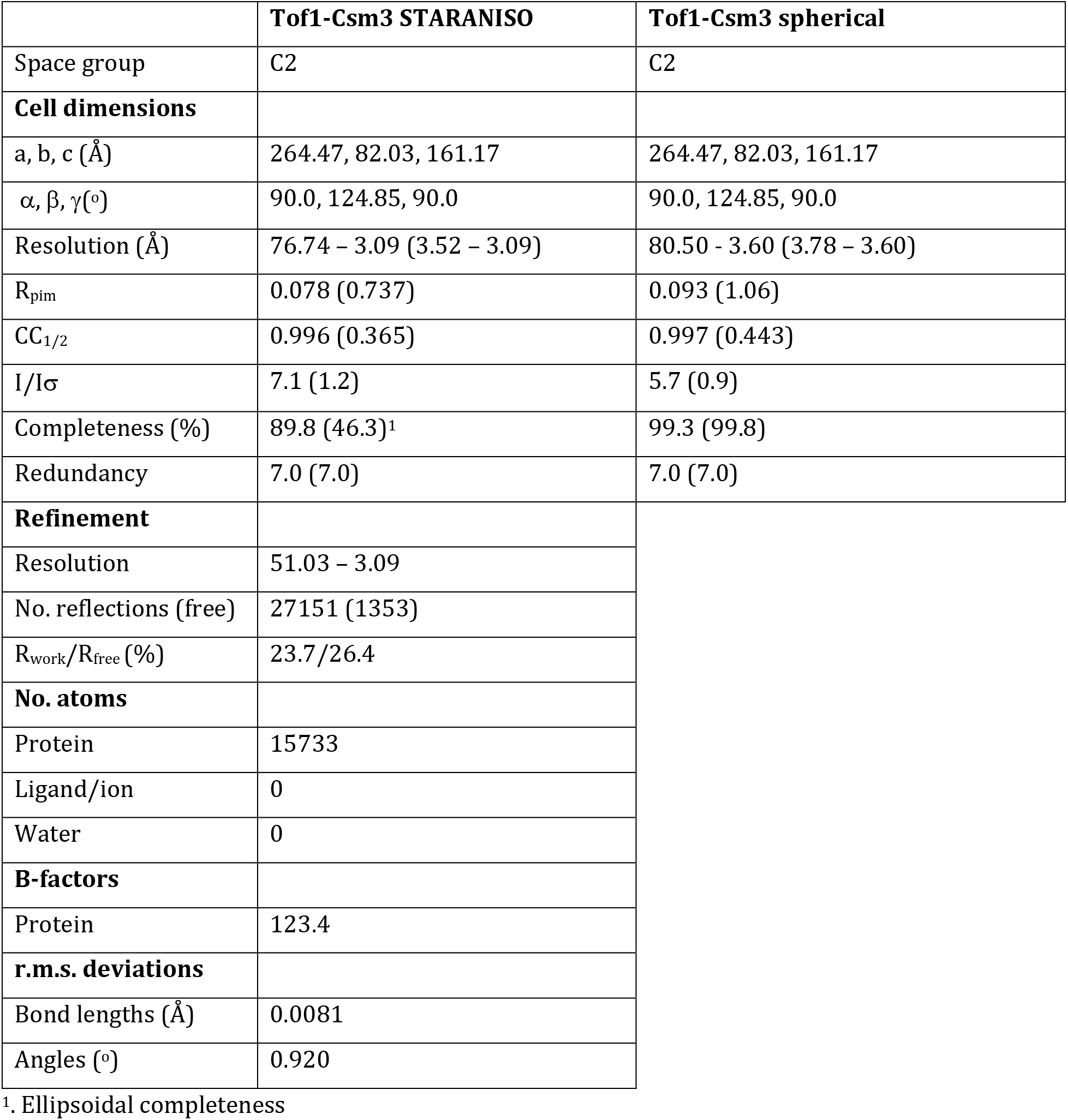
Crystallographic data collection and refinement statistics.

|  | **Tof1-Csm3 STARANISO** | **Tof1-Csm3 spherical** |
| --- | --- | --- |
| Space group | C2 | C2 |
| **Cell dimensions** |  |  |
| a, b, c (Å) | 264.47, 82.03, 161.17 | 264.47, 82.03, 161.17 |
| $\alpha, \beta, \gamma(^{\circ})$ | 90.0, 124.85, 90.0 | 90.0, 124.85, 90.0 |
| Resolution (Å) | 76.74 – 3.09 (3.52 – 3.09) | 80.50 – 3.60 (3.78 – 3.60) |
| R<sub>pim</sub> | 0.078 (0.737) | 0.093 (1.06) |
| CC<sub>1/2</sub> | 0.996 (0.365) | 0.997 (0.443) |
| I/Iσ | 7.1 (1.2) | 5.7 (0.9) |
| Completeness (%) | 89.8 (46.3)<sup>1</sup> | 99.3 (99.8) |
| Redundancy | 7.0 (7.0) | 7.0 (7.0) |
| **Refinement** |  |  |
| Resolution | 51.03 – 3.09 |  |
| No. reflections (free) | 27151 (1353) |  |
| R<sub>work</sub>/R<sub>free</sub> (%) | 23.7/26.4 |  |
| **No. atoms** |  |  |
| Protein | 15733 |  |
| Ligand/ion | 0 |  |
| Water | 0 |  |
| **B-factors** |  |  |
| Protein | 123.4 |  |
| **r.m.s. deviations** |  |  |
| Bond lengths (Å) | 0.0081 |  |
| Angles (°) | 0.920 |  |

<sup>1</sup>. Ellipsoidal completeness

The structure reveals that, instead of containing distinct domains, the entire complex forms an extended alpha-helical repeat protein (Fig 1A). Tof1 begins with two helices linked by a beta-hairpin, which is then followed by eight 3-helix armadillo repeats (Arm1-8). The previous Timeless N-terminal domain structure is a fragment of this structure ending after Arm-5. For this fragment, Tof1 and Timeless are clearly highly structurally related (Fig. 1B). Csm3 further extends the alpha-helical repeat structure by forming a five-helix bundle which packs on the C-terminus of Tof1. Csm3 shows some similarity to a tetra-helical bundle helix-turn-helix fold, but the first helix is broken into two (Fig. 1C). The closest structural homologue identified by PDBeFold [39] is the DNA binding domain of the small terminase from bacteriophage SF6 (Fig. 1C) [40]. However, in contrast to this domain, the helical bundle of Csm3 is flattened such that it no longer has a hydrophobic core and thus does not appear to be an independently folded structure and rather acts as a cap on the C-terminus of Tof1 (Fig.1C). The interface between the two is largely hydrophobic with some salt bridges, and consists of a large percentage of Csm3 (Fig. 1D), showing why both proteins are required to stabilize each other [41]. α25 and α26 from Arm-8 plus the following helix α27 contain all the interaction sites for Csm3 (Fig. 1D). Tof1 helices α25 and α26 pack predominantly against Csm3 α3. Tof1 helix α27 inserts into the concave structure of Csm3 making hydrophobic interactions with Csm3 helices α2 and α3.

**Fig. 1.**
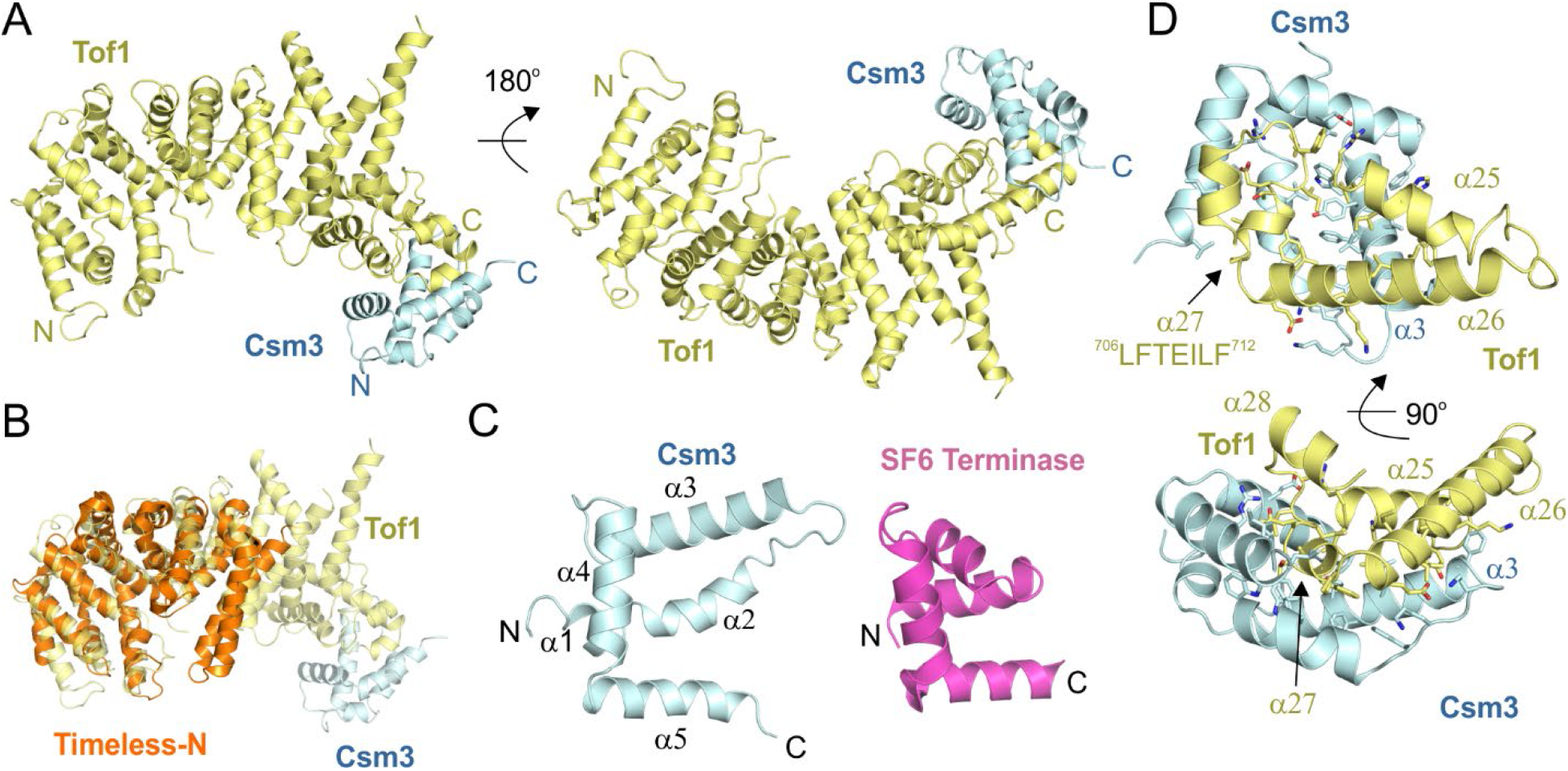
Crystal structure of the Tof1-Csm3 complex. (**A**) The complex is shown from two views with Tof1 colored in yellow and Csm3 in cyan. (**B**) Superposition of an N-terminal fragment of Timeless (PDB 5MQI) onto Tof1. (**C**) The structure of Csm3 and comparison to a protein with a similar fold, the DNA binding domain of SF6 small terminase (PDB 4ZC3). (**D**) Details of the interaction between Tof1 and Csm3. Only α25-27 of Tof1 are shown, and interfacial residues are shown in stick representation.

### Relation of Tof1 homologues

With the structure of the fungal Tof1-Csm3 complex, it is possible to gain insight into the structures of the orthologous protein Timeless, and its homologue, the circadian rhythm regulator drosophila Timeless (CR-Timeless) (Fig. 2). From this analysis it is immediately clear that the Timeless protein from varying eukaryotes has the same overall fold, with the hydrophobicity profile of all helices up to α26 conserved. Interestingly, helix α27, the major Csm3/Tipin interaction site, is very highly conserved in all the DNA replication Tof1/Timeless proteins but absent in the drosophila CR-Timeless. This is clearly indicative of the separate function of this protein. Furthermore, Loop 1 is highly conserved in the DNA replication Tof1/Timeless proteins (Fig. S3A), but has an entirely different sequence in CR-Timeless. This suggests Loop 1 has an important role in DNA replication.

**Fig. 2.**
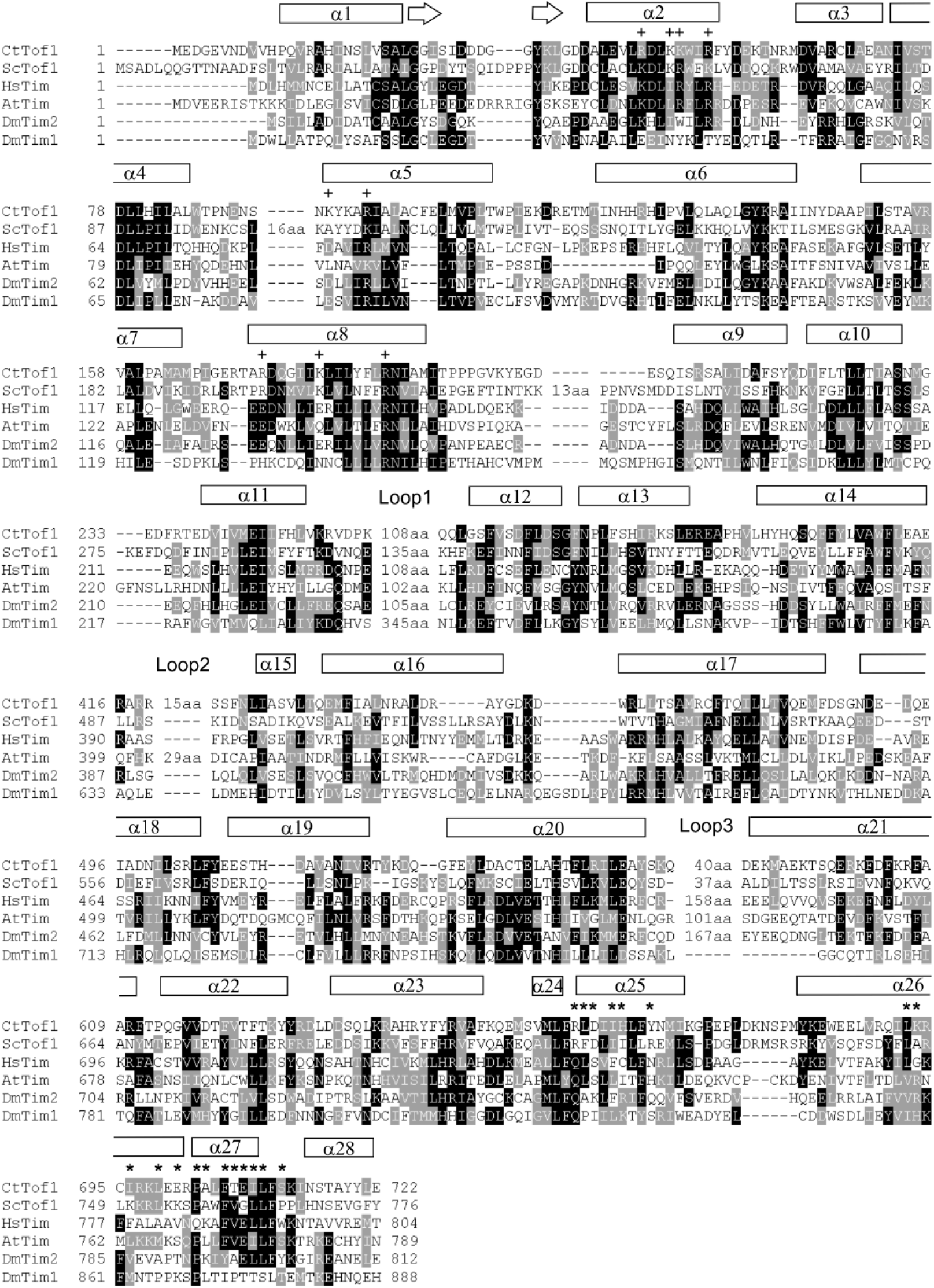
Sequence alignment of Tof1 with Timeless proteins. The alignment was performed using ClustalX2 [48] and displayed using BoxShade [49]. Ct – *Chaetomium thermophilum*, Sc – *Saccharomyces cerevisiae*, Hs – *Homo sapiens*, At – *Arabidopsis thaliana*, Dm – *Drosophila melanogaster*. Loop numbering refers to Fig S1A. **+** Peptide interacting, **\*** Csm3 interacting.

Interestingly, the presence of an additional small C-terminal portion of Tof1 (residues 728-900) largely increases the stability of the Tof1-Csm3 complex, despite preventing crystal diffraction (Fig. S1). The Swiss-model server identifies this as the human Timeless PARP1-binding PAB domain (Fig. S3B). This domain can also be found in the CR-Timeless sequence (Fig. S3B), but not *Arabidopsis* Timeless. Confirming this assignment, the RaptorX server, which determines the fold *de novo* by evolutionarily predicted sequence-contact restraints, independently generates a structure from the *C. thermophilum* sequence which is highly similar to the human PAB domain (Fig. S3C). This assignment is particularly notable because budding yeast contains no PARP proteins, and indeed important PARP1-binding residues are not conserved in the fungal proteins (Fig. S3B). Surprisingly, these residues are conserved in CR-Timeless.

### Interactions of the Tof1-Csm3 complex

Other Armadillo repeat proteins, such as β-catenin and importin-α, often bind peptides within the interior of their α-solenoid structure [42, 43]. As previously noted, there is a highly conserved cleft within this interior towards the N-terminus of Tof1 [34], lined with basic and hydrophobic residues. In our structure, the purification tag of a symmetry copy occupies this cleft (Fig. 3A). Furthermore, this interaction occurs in all three copies in the asymmetric unit despite the lack of symmetry in the packing, which suggests the patch has a very high propensity for peptide binding (Fig, 3A). Intriguingly, in human Timeless, one Arginine lining this pocket, Arg40 (*C. thermophilum* Lys51), has been found mutated six times to cysteine and once to proline in different cancers in the COSMIC Sanger database [44], This is the most common cancer-associated missense mutation of Timeless, with the second being Pro1043 (five times), which is a key Parp1-interacting residue (Fig. S3B). This implies Arg40 has functional importance. Additionally, some residues, such as Arg47 and Arg54, are highly conserved in fork protection Tof1/Timeless but not CR-Timeless (Fig. 2B). Considering the positive charge of this pocket we tested if this was a functional DNA binding site. In an EMSA, Tof1(long)-Csm3 bound to dsDNA but not ssDNA with a mild affinity (Fig. 3B). However, a quadruple alanine mutation of the peptide-binding patch (K50/R54/R98/R173) had no effect on this activity (Fig. 3B), suggesting that this patch has another function. As this N-terminal site is not involved in DNA binding, we next tested whether the other positively charged region of the complex harbored a DNA binding site – the C-terminus of Tof1 with Csm3 (Fig. 3C). We were able to generate a stable minimal complex of Tof1-Csm3 containing only helices α22-28 of Tof1 with Csm3 (Fig S1A). Interestingly, this minimal complex retained dsDNA binding activity (Fig. 3B), although *circa* 2-fold weaker, suggesting this region of the complex contains a significant portion of the dsDNA binding site.

**Fig. 3.**
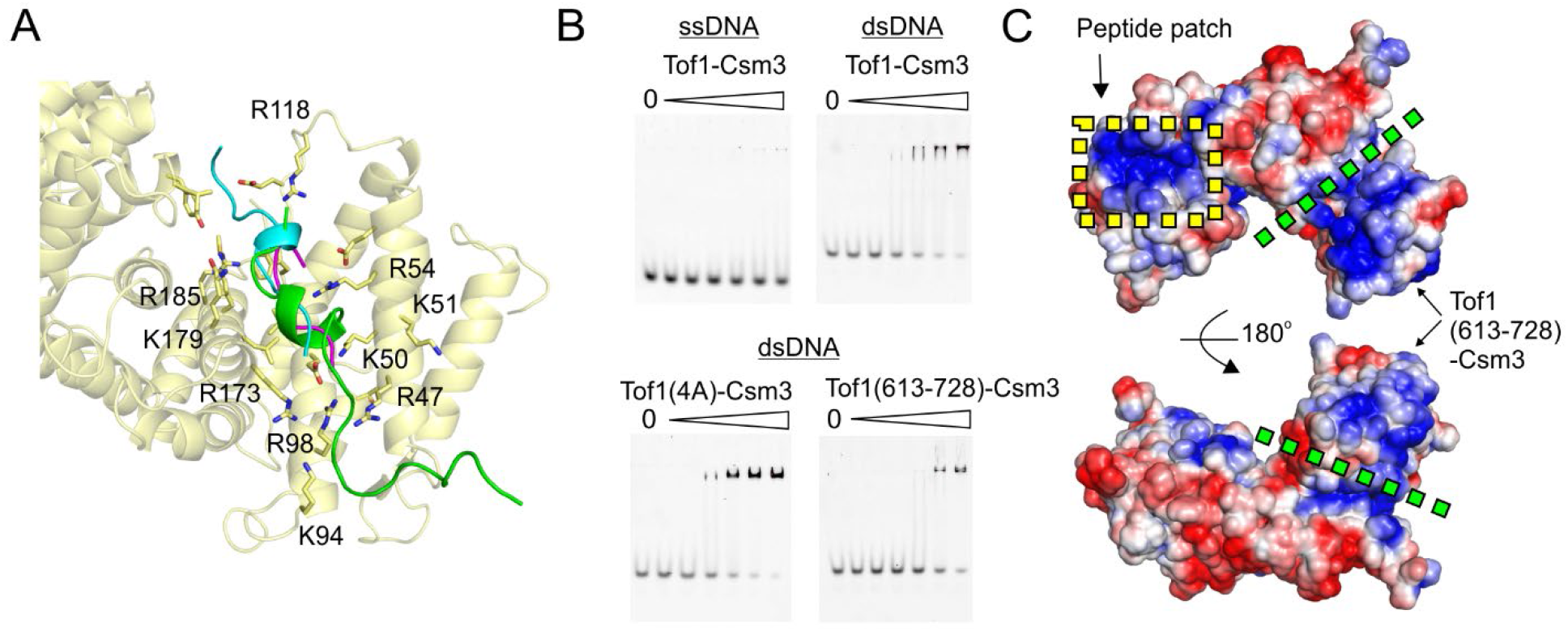
Interactions of the Tof1-Csm3 complex. (**A**) Interaction between Tof1 and a symmetry-related purification tag. The peptide interacting with each copy of Tof1 was superimposed by aligning the interacting elements of Tof1, and is shown in cartoon representation. The N-terminus of Chain E (interacting with Tof1 chain C) is in green, the N-terminus of Chain A (interacting with Chain E) is in cyan, and the N-terminus of Chain D (interacting with Chain A) is magenta. (**B**) DNA binding properties of the Tof1-Csm3 complex. The indicated complex was added in a 2-fold dilution series from 4 μM to 0.125 μM. (**C**) Surface potential of the Tof1-Csm3 complex. The potential was calculated using APBS [50]. Positively charged surface is colored blue and negatively colored red. The green dotted line indicates the approximate position of the indicated Tof1 truncation.

## Discussion

The Tof1-Csm3 complex and its human orthologue the Timeless-Tipin complex have been shown to have a number of important functions at the replication fork. By solving the structure of this complex we now can begin to understand what molecular role the complex plays to link these processes together. Importantly, we show that the Tof1-Csm3 and Timeless-Tipin complexes are highly related (Fig. 1B, 2), and thus likely play the same role at the fork, while the Drosophila CR-Timeless has the same overall fold but clear differences in sequence related to its separate role (Fig. 2) [31], i.e. interaction with the Period [45] and Cryptochrome proteins [46].

Rather than consisting of multiple distinct domains, the core of the Tof1-Csm3 complex forms a single large alpha-helical repeat protein. Thus, some caution should be taken with any previous functional studies which artificially truncated this fold. This structure suggests the protein has a scaffolding or mechanical role at the fork, which is presumably further regulated by the large intrinsically disordered regions in both Tof1 and Csm3. A recent 2D cryo-electron microscopy study has shown that Tof1-Csm3 binds ahead of the replicative helicase and reduces the flexibility of incoming DNA [35]. Such a mechanical function would fit well with my structure, given its rigid structure and dsDNA binding properties. My structure also reveals a highly conserved peptide-binding patch that may interact with a partner protein such as the replicative helicase [14] or Mrc1/Claspin [11, 47]. Intriguingly one arginine in this binding patch has been detected as a cancer-associated mutation. Normally, Timeless is overexpressed in cancer, and cells become dependent on the protein to combat cancer-caused replication stress [6]. Future studies will be able to ascertain whether this mutation has a positive or negative effect on Timeless activity.

Overall, our structure provides a basis for understanding the interactions, mutations and function of both the fork protection complex and circadian rhythm regulator Timeless protein at a molecular level.

## Acknowledgements

I would like to thank beamline scientists and support staff at the EMBL-operated PETRA III beamline P14 at DESY. This work was funded by DFG grant GR5152/3-1 and a postdoctoral fellowship from the Alexander von Humboldt Foundation.

## Data Availability

Coordinates and structure factors for the Tof1-Csm3 complex have been deposited in the protein data bank under the PDB code 6XWX.

## Methods

### Molecular biology

The entire *Chaetomium thermophilum TOF1* and *CSM3* genes were synthesized by ATG:biosynthetics and provided in separate pGH vectors. For all expression constructs a pETM-14 vector (EMBL) with an N-terminal His6-tag. *TOF1* and *CSM3* were cloned into the multi-cloning site for co-transcription and Csm3 was untagged but cloned in with a ribosomal binding sequence. Construct design was guided by disorder prediction by the DISOPRED server [51]. First Tof1(1-728) was amplified with primers 5’CAAGA<u>CCCATGG</u>AAGATGGTGAGGTTAACGATG and 5’CAAGAC<u>GGATCC</u>TTATTGTTTCTCAAAGCCGTATTCCAG, and inserted by restriction cloning into pETM-14 using NcoI and BamH1 to generate pETM-14-Tof1. Csm3(48-157) or Csm3(77-157) was then amplified with 5’ CAAGAC<u>AAGCTT</u>TTATTCAAATGATGCTTTCGGGCGGAG and either 5’CAAGAC<u>GAATTC</u>AAGAAGGAGATATACCATGACAGATGCGCTGGGTATCGAC or 5’CAAGAC<u>GAATTC</u>AAGAAGGAGATATACCATGTCGGAAAAAGGTATTCCTAAACTTCG to generate either pETM-14-Tof1-Csm3 or pETM14-Tof1-Csm3trunc. Loops were deleted from Tof1 by blunt-end mutagenesis. For DL1, residues 256-363 were replaced by one glycine using primers 5’<u>GT</u>CAGCAGCTTGGTAGCTTTGTTTCAG and 5’<u>C</u>TTTAGGGTCCACGCGTTTCACC. For DL1a, a construct containing DL1 was further mutated using primers 5’<u>GCACGC</u>CAGCAGCTTGGTAGCTTTGTTTCAG and 5’<u>GCGTTC</u>ACCTTTAGGGTCCACGCGTTTCAC resulting in replacement of residues 256-259 by a single glycine. For DL2, residues 420-434 were removed with primers 5’TCCAGTTTTAATTTAATTGCCTCCGTGC and 5’ACGGCGCGCGCGTTCGG. For DL3, residues 558-585 were replaced by one glycine using primers 5’<u>GT</u>TCGGCGGACGATGAAAAAATGG and 5’<u>C</u>GGATCGCACCTGAAGGTCTACG. The C-terminal domain of Tof1 (residues 728-900) was cloned into pETM14 with restriction enzymes BamHI and EcoRI using primers 5’CAAGAC<u>GGATCC</u>ACTGTTTCATCTAATCCACGTCCG and 5’CAAGAC<u>GAATTC</u>TTATTTGCGGCGCAATTGATGTTCAGC. The same approach was used to C-terminally elongate Tof1-Csm3. In this case, the stop codon and BamHI site were then removed by blunt-end mutagenesis using primers 5’ACTGTTTCATCTAATCCACGTCCG and 5’TTGTTTCTCAAAGCCGTATTCCAG to generate pETM14-Tof1long-Csm3. For DL4, residues 784-813 were replaced by a single glycine using primers 5’<u>GT</u>GCATACACCACCGTTCGCCC and 5’<u>C</u>CGCTCGACGTTCGCGTTCCG. To generate Tof1-Csm3 constructs where residues 10-612 of Tof1 were removed, blunt-end mutagenesis was performed with primers 5’ACCCCACAAGGCGTGGTAGATACG and 5’TACATCGTTAACCTCACCATCTTCC using either pETM14-Tof1-Csm3 or pETM14-Tof1long-Csm3 as templates to generate pETM14-NtruncTof1-Csm3 and pETM14-NtruncTof1long-Csm3. The quadruple alanine substitution was generated by sequential blunt-end mutagenesis using primers: K50A – 5’AAAATGGATTCGTTTCTATGACGAAAAAACTAACC and 5’GCCAGGTCCCGAAGGACTTCC, R54A – 5’TTTCTATGACGAAAAAACTAACCGCATGG and 5’GCAATCCATTTTGCCAGGTCCCGAAGG, R98A – 5’CTATC GCATTGGCGTGCTTCGAACTG and 5’CCGC TTTGTATTTATTCGAGTTTTCATTCG, R173A – 5’CCGACCAAGGCATCATAAAACTGATTCTG and 5’CTGCGGTTCGTTCTCCGATCGG

### Expression and purification of Tof1-Csm3 constructs

All Tof1-Csm3 constructs were expressed and purified using the same method. The appropriate plasmid was transformed into BL21(DE3)Star (Novagen) cells and grown in media supplemented with 50 μg/ml kanamycin. Large terrific broth expression cultures were inoculated 1 in 100 with an overnight start culture and grown at 30°C with shaking at 200 rpm until they reached an A_600_ of 0.5. The temperature was reduced to 17°C, and cultures were induced with 0.4 mM IPTG overnight, followed by harvesting using centrifugation, and storage of bacterial pellets at −80°C until use. Thawed pellets were resuspended in 50 mM Tris-HCl pH 8.0, 500 mM NaCl, 1 mM TCEP, 10 mM imidazole, 1 EDTA-free protease inhibitor tablet/50 mL (Roche), and 30 U/mL DNase I. Lysis was performed by two passages through a Microfluidics M-110P microfluidizer at 150 MPa. The lysate was cleared by centrifugation for 1 hour at 60000 x g and then loaded on a 5 mL Histrap FF column (GE Healthcare) equilibrated in 50 mM Tris-HCl pH 8.0, 500 mM NaCl, 1 mM TCEP, using an Akta Purifier FPLC system. The column was then washed with 16 column volumes of 25 mM imidazole in the same buffer, and then eluted with a 10 column volume gradient of 25 – 250 mM imidazole. Protein-containing fractions were then concentrated by microfiltration before loading on a Superdex 200 26/60 prepgrade column (GE Healthcare) column that had been equilibrated in 20 mM HEPES pH 7.5, 1 mM TCEP with different concentrations of NaCl for the different constructs as indicated in Figure S1A (HBS-X). Fractions containing the Tof1-Csm3 complex were then concentrated using a spin concentrator and stored at −80°C. The protein concentration was estimated from the A_280_ absorption using the extinction coefficient calculated by the Expasy Protparam server [52].

### Crystallization

Crystallization trials were performed with a Honeybee fluid transfer robot using the sitting drop vapor-diffusion method with 0.3 μL of protein was mixed at a 1:1 ratio with mother liquor from customized screen (Fig. S1B). Drops were incubated at 20°C and equilibrated against 40 μL mother liquor supplemented with an additional 0, 125, 187 or 250 mM NaCl. The Tof1-Csm3 complex was crystallized at 30 mg/ml in HBS-500 and mixed with a precipitant solution containing 0.2 M Potassium Acetate, 4-8% PEG 20000, 0.1 M Tris-HCl pH 7.5-8.5. The reservoir was supplemented with 125 mM NaCl. Crystals were cryo-protected in the same solution with 25% ethylene glycol additional.

### Data collection, Structure solution and refinement

The data were collected at 100K and at a wavelength of 0.980 Å at beamline P14 of the EMBL-operated PETRA III ring at DESY. The data were integrated using XDS [53], and then merged either using Aimless [54] to 3.6 Å or using the STARANISO server [55] to 3.09 Å (Table 1).

A homology model of *C. thermophilum* Tof1 residues 1-488 was generated from the structure of the corresponding region from human timeless [34] (PDB 5MQI) using the SWISS-MODEL server [56]. From this, all sidechains were truncated to alanine and all loops deleted. Three copies were found using the STARANISO-processed data and PHASER [57] with a TFZ score of 11.1. Initial refinement was performed using Phenix Refine [58] and the non-corrected data to 3.6 Å. Helices were manually placed in the density, and Phenix Autobuild [59] was run occasionally and reduced model bias. A continuation of the alpha-helical repeat structure from the Timeless fragment was clear, and so this was exploited for sequence and topology assignment. The sequence was too short for the last four helices and so these were assigned to Csm3. Once the model was largely complete, refinement was continued using Buster [60, 61] and the anisotropically corrected data to 3.1 Å. Intermittent cycles of molecular dynamic force-field refinement using the Namdinator server [37] proved essential for overcoming model bias, and the RaptorX evolutionary contact server [38] was used to validate the sequence assignment. The final R_work_/R_free_ values were 23.7/26.4% with 0.21% Ramachandran outliers, a Clashscore of 2.80 and an overall MolProbity score of 1.84 [62]. The Chain A-Chain B Tof1-Csm3 copy was the best defined and used for all structural figures, unless otherwise stated, and these were generated using PyMOL [63].

### Electrophoretic Mobility Shift Assays

The ssDNA oligonucleotide had the following sequence: 5’-Cy3-GTAGTTTGTACTGGTGACGA. The dsDNA substrate was generated by mixing this 1:1 with the complementary oligonucleotide 5’-TCGTCACCAGTACAAACTAC, melting at 95°C for 2 minutes, and then slow cooling at room temperature to anneal. DNA substrates were used at a final concentration of 50 nM. Samples were prepared in HBS-200 with an additional 10% glycerol, and incubated on ice for 30 minutes. Samples were loaded on a 6% polyacrylamide gel and run at 70 V for 50 minutes in a tris-glycine buffer system. After running, gels were scanned with a Pharos FX fluorescence imaging system (Biorad) and excitation/emission wavelengths of 532/605 nm.

