## Supplementary Information for "Crystal structure of the Tof1-Csm3 (Timeless-Tipin) fork protection complex"

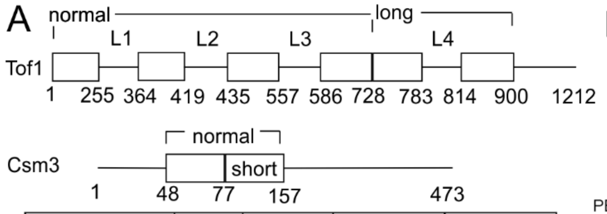


| Construct | NaCl (mM) | Crystals | Diffraction | Solved |
|---|---|---|---|---|
| Tof1-Csm3 | 500 | - | - | - |
| Tof1-Csm3 DL1/DL3 | 500 | Yes | V. weak | - |
| Tof1-Csm3 DL1a/DL3 | 500 | No | - | - |
| Tof1-Csm3short DL1/DL3 | 500 | Yes | No | - |
| Tof1long-Csm3 DL1/DL3 | 200 | No | - | - |
| Tof1long-Csm3 DL1/DL3/DL4 | 200 | Yes | No | - |
| Tof1-Csm3 DL1/DL2/DL3 | 500 | Yes | 3.1 A | Yes |
| Tof1-Csm3short DL1/DL2/DL3 | 500 | Yes | No | - |
| Tof1-C | 0 | Yes | 2.9 A | No – Severe crystal pathologies |
| Tof1-C DL4 | 0 | No | - | - |
| NtruncTof1-Csm3 | 200 | Yes | No | - |
| NtruncTof1long-Csm3 | 50 | Yes | No | - |
| NtruncTof1long-Csm3 DL4 | 50 | Yes | No | - |

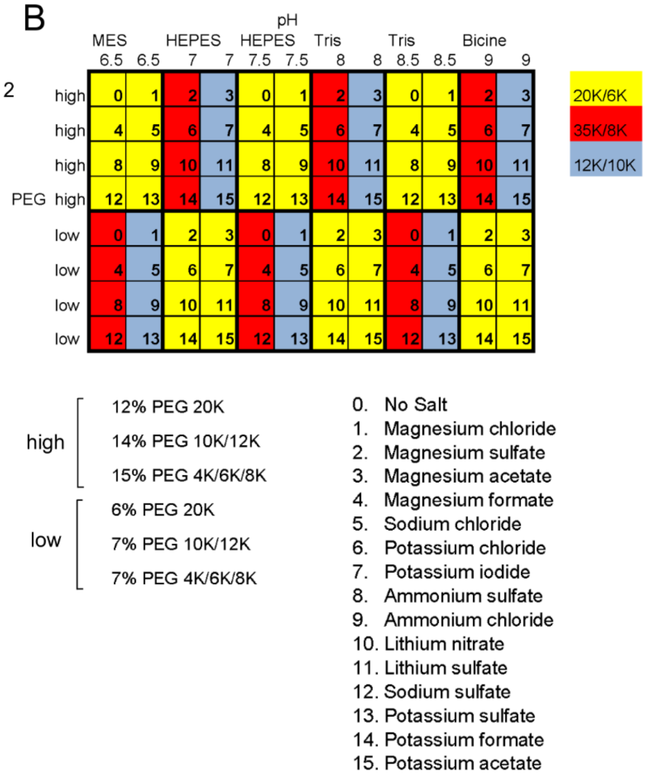


**Figure S1 – Crystallization strategy.** (**A**) Constructs screened for crystallization. 'Short' contains only the indicated region, while 'long' contains the indicated region additional. Tof1-C only contains the 'long' region. 'NaCl' refers to the concentration of sodium chloride required to solubilize the complex. (**B**) Design of homemade crystallization screens for challenging complexes. One screen contains PEGs 4K, 6K and 8K, while the other contains 10K, 12K and 20K. All salt is used at a concentration of 200 mM. The salts (numbered 0-15) and the high and low PEG concentrations are described below the schematic.

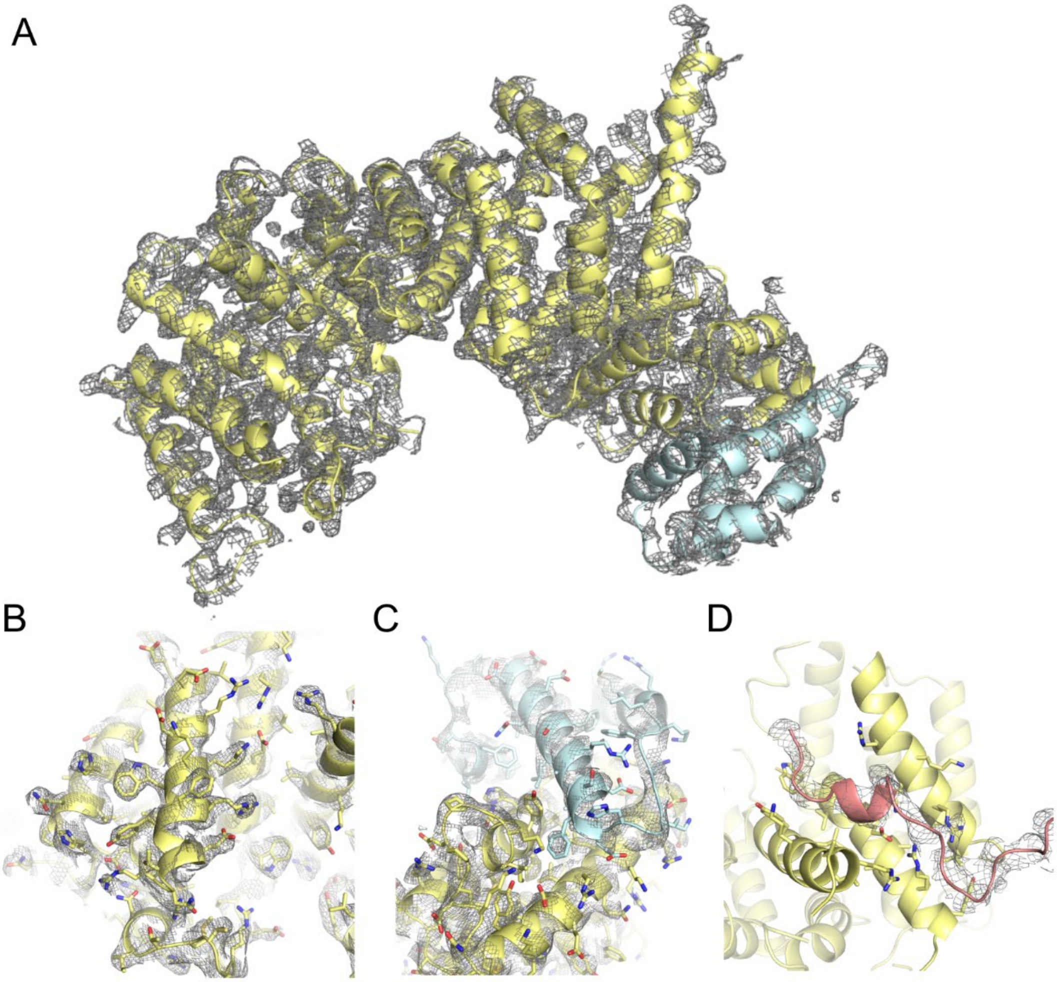


**Figure S2 – Quality of the electron density and model fit.** Models are shown in cartoon and stick representation with the same coloring as Fig. 1. 2Fo-Fc maps are shown for (**A**) The entirety of the Tof1-Csm3 complex (Chain A-Chain B), (**B**) A low B-factor region of Tof1, (**C**) The interface between Tof1 and Csm3, (**D**) The N-terminal purification tag of Chain E (interacting with Tof1 chain C).

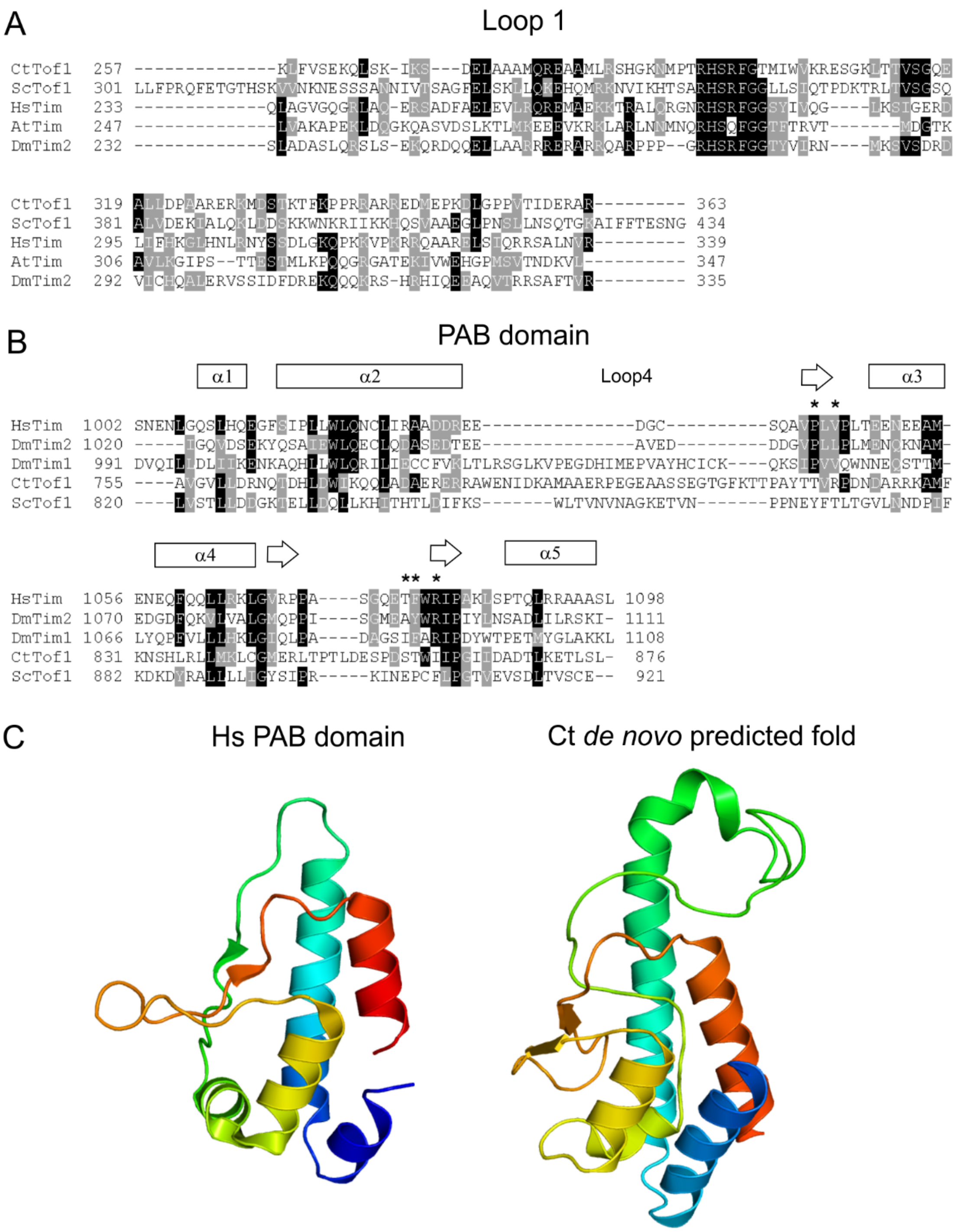


**Figure S3 – Sequence analysis of parts of the Tof1-Csm3 complex which were not crystallized.** Alignments were prepared as in Fig. 2. (**A**) Alignment of Loop 1 (Fig. S1A) from Tof1/Timeless homologues, excluding CR-Timeless, as the sequence and size are very different. (**B**) Alignment of the putative PAB domains from Tof1/Timeless homologues. Arabidopsis Tof1 was excluded as no PAB domain is predicted to be present. Loop4 is shown in Fig. S1A. (**C**) Comparison of the fold of the Timeless PAB domain (PDB 4XHT) with the structure of the C-terminal sequence of Chaetomium Tof1 generated *de novo* by the RaptorX server (Wang *et al.*, 2017)
